# Inhibition of Casein Kinase 2 induces cell death in chronic myelogenous leukemia cells resistant to tyrosine kinase inhibitors

**DOI:** 10.1101/2021.06.28.450156

**Authors:** Ondřej Mitrovský, Denisa Myslivcová, Tereza Macháčková-Lopotová, Adam Obr, Kamila Čermáková, Šárka Ransdorfová, Jana Březinová, Hana Klamová, Markéta Žáčková

## Abstract

Chronic myelogenous leukemia (CML) is a myeloproliferative disease characterized by the presence of a BCR-ABL oncogene. Despite the high performance of treatment with tyrosine kinase inhibitors (TKI), about 30 % of patients develop resistance to the therapy. To improve the outcomes, identification of new targets of treatment is needed. Here, we explored the Casein Kinase 2 (CK2) as a potential target for CML therapy. Previously, we detected increased phosphorylation of HSP90β Serine 226 in patients non-responding to TKIs imatinib and dasatinib. This site is known to be phosphorylated by CK2, which was also linked to CML resistance to imatinib. In the present work, we established six novel imatinib- and dasatinib-resistant CML cell lines, all of which had increased CK2 activation. A CK2 inhibitor, CX-4945, induced cell death of CML cells in both parental and resistant cell lines. In some cases, CK2 inhibition also potentiated the effects of TKI on the cell metabolic activity. No effects of CK2 inhibition were observed in normal mononuclear blood cells from healthy donors and BCR-ABL negative HL60 cell line. Our data indicate that CK2 kinase supports CML cell viability even in cells with different mechanisms of resistance to TKI, and thus represents a potential target for treatment.

## Introduction

Chronic myeloid leukemia (CML) is a myeloproliferative disease characterized by the presence of Philadelphia translocation t(9;22)(q34;q11), resulting in the expression of a constitutively active BCR-ABL kinase. BCR-ABL is the main driver of CML (1). A selective BCR-ABL inhibitor, imatinib, has been introduced into clinical practice in 2001 and represents a breakthrough in CML therapy (2,3). About one-third of patients, however, develop resistance or intolerance to the drug. To address this issue, second- and third-generation tyrosine kinase inhibitors (TKIs) — like dasatinib, nilotinib, and ponatinib — were developed, and are effective against most imatinib-resistant cases of CML.

Patients treated with advanced generations of TKIs, however, also have a high risk of acquiring resistance to the new drugs (4–6). Therefore, other ways of targeting resistant CML cells are explored. The research in this area focuses on strategies to target pathways that are not induced by BCR-ABL, including those currently considered as non-oncogenic (7,8).

Casein kinase 2 (CK2) is a ubiquitously expressed serine/threonine (S/T) kinase. It is usually present as a tetrameric complex of two catalytic (α, α’) and two regulatory (β) subunits. Although it was previously labeled as constitutively active, several reports now suggest many possible pathways of CK2 regulation, e.g. via protein interactions, differential phosphorylation, and interactions with other regulatory elements (9). CK2 has a net pro-survival, anti-apoptotic role (10–13). Its expression is increased in cancer cells (14–16), and its signaling feeds the usual cancer-related pathways, even reaching the point where cancer cells become CK2-dependent (17).

Furthermore, CK2 has also been associated with multi-drug resistance to chemotherapy (18,19). This all shows CK2 as an important player in cancer pathogenesis, and since some highly specific inhibitors are available, it also represents a possible target for treatment (9). An orally available compound Silmitasertib (CX-4945) is currently evaluated in clinical trials for several cancer types (20), but so far has not been tested for CML treatment (21).

We used protein-antibody arrays to find proteins differentially expressed and/or phosphorylated in samples of patients divided into groups according to their therapy response (as characterized by ELN (22)), and healthy donors. Among others, we detected increased phosphorylation of HSP90β on Ser226 in imatinib- and dasatinib-resistant patients. Phosphorylation of this site was previously identified as CK2-dependent (23,24). As CK2 is associated with resistance to imatinib (25,26), we further explored the role of CK2 in the resistance to both imatinib and dasatinib. In TKI-resistant cells, CK2 expression was increased, and all the resistants were also highly sensitive to CK2 inhibition. Additionally, our preliminary data from patients non-responding to the therapy showed corresponding effects of CK2 inhibition on cell viability.

## Results

### Phosphorylation of HSP90β S226, a CK2-target, is increased in patients non-responding to TKI

Using protein-antibody arrays, we analyzed 14 samples of total leukocytes obtained from CML patients with different responses to the standard therapy (supplementary figure 1, table 1A). Several proteins were differentially phosphorylated in patient samples compared to healthy donors. Besides the usual suspects, such as BCR-ABL and SFK, we found changes at phosphorylation sites of the heat-shock proteins HSP90 and HSP27 (supplementary figure 1). We identified a systematic increase of HSP90β S226 phosphorylation in samples from patients non-responding to therapy, or relapsed, compared to healthy donors. For two another patients, we performed the screen at the time of diagnosis and subsequently 10 and 11 months after treatment initiation (table 1B). The ratio of HSP90β phosphorylated on S226 to total HSP90β was markedly higher in the non-responding patient.

**Table 1A.** Patient characteristics at the time of analysis. HD – healthy donors, OR – optimal response to treatment, NR – nonresponding patients, REL – relapsed patients. DG – diagnosis, TH – therapy (imatinib/dasatinib/nilotinib), BCR-ABL (%) – mRNA level, WBC – white blood cell count, PLT – platelet count, BL % of blast cells in total cell count, mut – mutations in the BCR-ABL kinase domain, ab – karyotype aberations

| Tab 1A | F/M | months from DG | TH (I/D/N) | BCR-ABL (%) | WBC (10<sup>9</sup>/l) | PLT (10<sup>9</sup>/l) | BL (%) | mut (n) | ab (n) |
| --- | --- | --- | --- | --- | --- | --- | --- | --- | --- |
| OR (n=4) | 4/0 | 70,3 (12-141) | 3/0/1 | 0 (0-0,001) | 5,35 (3,98-14,4) | 202 (170-268) | 0 | 0 | 0 |
| NR (n=5) | 3/2 | 52,9 (12-141) | 3/2/0 | 13 (4,7-33) | 4,3 (3,26-7,79) | 198 (95-212) | 0 | 1 | 1 |
| REL (n=5) | 3/2 | 52 (33-164) | 2/2/1 | 57,5 (1,1-109) | 10,88 (3,9-22,05) | 204 (52-420) | 0 | 2 | 2 |
| HD (n=5) | 3/2 | N/A | N/A | N/A | N/A | N/A | N/A | N/A | N/A |

**Table 1B.** Patient characteristics at the time of analysis. NR – nonresponding patient, OR – optimal response to treatment, DG – diagnosis, BCR-ABL (%) – mRNA level, WBC – white blood cell count, PLT – platelet count, BL % of blast cells in total cell count, TH – therapy

| Tab 1B | sex | age | months from DG | BCR-ABL (%) | WBC (10<sup>9</sup>/l) | PLT (10<sup>9</sup>/l) | BL (%) | TH |
| --- | --- | --- | --- | --- | --- | --- | --- | --- |
| NR | F | 49 | 0 | 18,4 | 66,59 | 454 | 1 |  |
|  |  |  | 11 | 4,7 | 5,75 | 178 | 0 | imatinib |
| OR | F | 67 | 0 | 52 | 56,11 | 502 | 1 |  |
|  |  |  | 11 | 0,001 | 5,91 | 216 | 0 | imatinib |

### Establishment and characterization of six novel BCR-ABL positive cell lines resistant either to imatinib or dasatinib

We established six new TKI-resistant, CML-derived cell lines. JURL-MK1, MOLM-7, and K562 cells able to grow in media containing 2 μM imatinib or 2 nM dasatinib, respectively, are further denoted as “TKI resistant” (imatinib resistant – IR, dasatinib resistant – DR). BCR-ABL negative cell lines HL-60 and OCI-AML3 were cultivated under the same conditions as CML cell lines and were used as controls. The properties of the new cell lines were characterized as described below.

### Cell growth

Cell growth and viability were assessed by trypan blue staining after treatment with either 10 μM imatinib or 100 nM dasatinib (Supplementary Figure S2). All the resistant sub-lines demonstrated decreased sensitivity to TKIs, compared to their parental cell lines. Notably, various levels of cross-resistance (i.e. resistance to both TKIs used) were detectable in all the resistant cells. The growth of BCR-ABL negative cells remained unaffected by the presence of TKIs.

### Sensitivity to TKI

Table 2 summarizes the EC50 values calculated from the measurement of cell activity after 48 h TKI treatment (Supplementary figure S2). All the resistant sub-lines exhibited lower sensitivity / higher EC50 values to both TKIs. Similar to the results of cell growth experiments, various levels of cross-resistance were evident in all the sub-lines.

**Table 2.** EC50 values of imatinib and dasatinib effects on the proliferation of sensitive and resistant cells. EC50 values of imatinib and dasatinib on all used cells. Cell lines were incubated with 0-100 μM imatinib or 0-100 nM dasatinib. The EC50 on cell proliferation/viability was assessed by the Alamar Blue. EC50 values and CI95% were calculated from 2-3 independent experiments.

| | imatinib (μM) | dasatinib (nM) |
| --- | --- | --- |
| **JURL-MK1** | **0,18** (0,16-0,20) | **0,16** (0,14-0,19) |
| **JURL-MK1 IR** | **25,3** (17,2-38,1) | **35,7** (23,5-63,3) |
| **JURL-MK1 DR** | **0,7** (0,6-0,9) | **2,0** (1,5-2,6) |
| **MOLM-7** | **0,25** (0,19-0,31) | **0,15** (0,11-0,19) |
| **MOLM-7 IR** | **59,2** (12,6-188,0) | **1,4** (0,7-2,6) |
| **MOLM-7 DR** | **92,7** (35,7-783,8) | **36,7** (26,3-55,7) |
| **K562** | **0,25** (0,12-0,42) | **0,34** (0,25-0,47) |
| **K562 IR** | **4,8** (3,5-6,5) | **3,5** (1,8-6,8) |
| **K562 DR** | **5,2** (2,9-9,3) | **3,6** (1,7-7,5) |

### BCR-ABL transcript variants and mutations in the BCR-ABL kinase domain

To further characterize the novel cell lines, we verified the type of BCR-ABL transcript present in the cells. The variant found in the parental line was always preserved in the derived resistant sub-lines. As MOLM-7 cells contained the e13a2 (b2a2) transcript, in JURL-MK1 and K562 the e14a2 (b3a2) transcript was detected.

In both JURL-MK1 IR and DR cells, there was a T315I mutation in the kinase domain. In resistant cells derived from MOLM-7 and K562, respectively, no mutation in the kinase domain of BCR-ABL was detected.

### Cytogenetics

Detailed results of cytogenetic verification of the new cell lines are shown in Supplementary Figure S3.

In all JURL-MK1 derived cells, chromosomal aberrations were almost identical to their parental line. Only polyploid mitoses were present, and the BCR-ABL fusion signal was detected on the same chromosomes in all the three JURL-MK1 cell lines (Supplementary Figure S3A).

In TKI-sensitive MOLM-7 cells (Supplementary Figure S3B), we detected structural changes relating to the BCR-ABL fusion gene. The control cells had two extra numerical Ph chromosomes. MOLM-7 IR cells were shown to be polyploid, and all BCR-ABL signals were present in two copies. One extra numerical Ph was present, and a second one was translocated and amplified on the short arm of a derivative chromosome 13. In MOLM-7 DR cells, again, one extra numerical Ph chromosome was detected, and a second one was translocated to the short arm of chromosome 18. Additionally, in all MOLM-7 cells, the BCR-ABL fusion signal was also detected on the short arm of chromosome 6.

The K562 cells used in this study are near triploid and posess characteristic markers as previously reported (27,28). One copy of the chromosomes X, 3, 9, 13, and 14 was lost in all K562 cells. In all the three K562 sub-lines the same abnormalities were present on several chromosomes (Supplementary Figure S3C). Additionally, some differences were detected. Results of fluorescence in-situ hybridization (FISH) with BCR-ABL specific probes showed that K562 and K562 DR (but not K562 IR) had BCR-ABL fusion on the short arm of the derivative chromosome 1. In K562 and K562 IR (but not K562 DR), this aberration was also found on the long arm of the derivative chromosome 22. In K562 IR, other non-amplified BCR-ABL fusions were detected on the long arms of the derivative chromosomes 2 and 5. In K562 DR, an additional BCR-ABL amplified signal was detected on the long arm of the derivative chromosome 11, and a non-amplified signal in the long arm of chromosome 16 and on the short arm of the chromosome X.

### BCR-ABL activity

The activity of BCR-ABL was assessed via evaluating the phosphorylation status of CRKL, a surrogate marker of BCR-ABL activity. In JURL-MK1 and MOLM-7 TKI-resistant sub-lines, P-CRKL levels were increased compared to parental cells. In K562 IR cells, interestingly, CRKL phosphorylation was significantly decreased compared to their sensitive counterparts (Figure 1).

**Figure 1.**
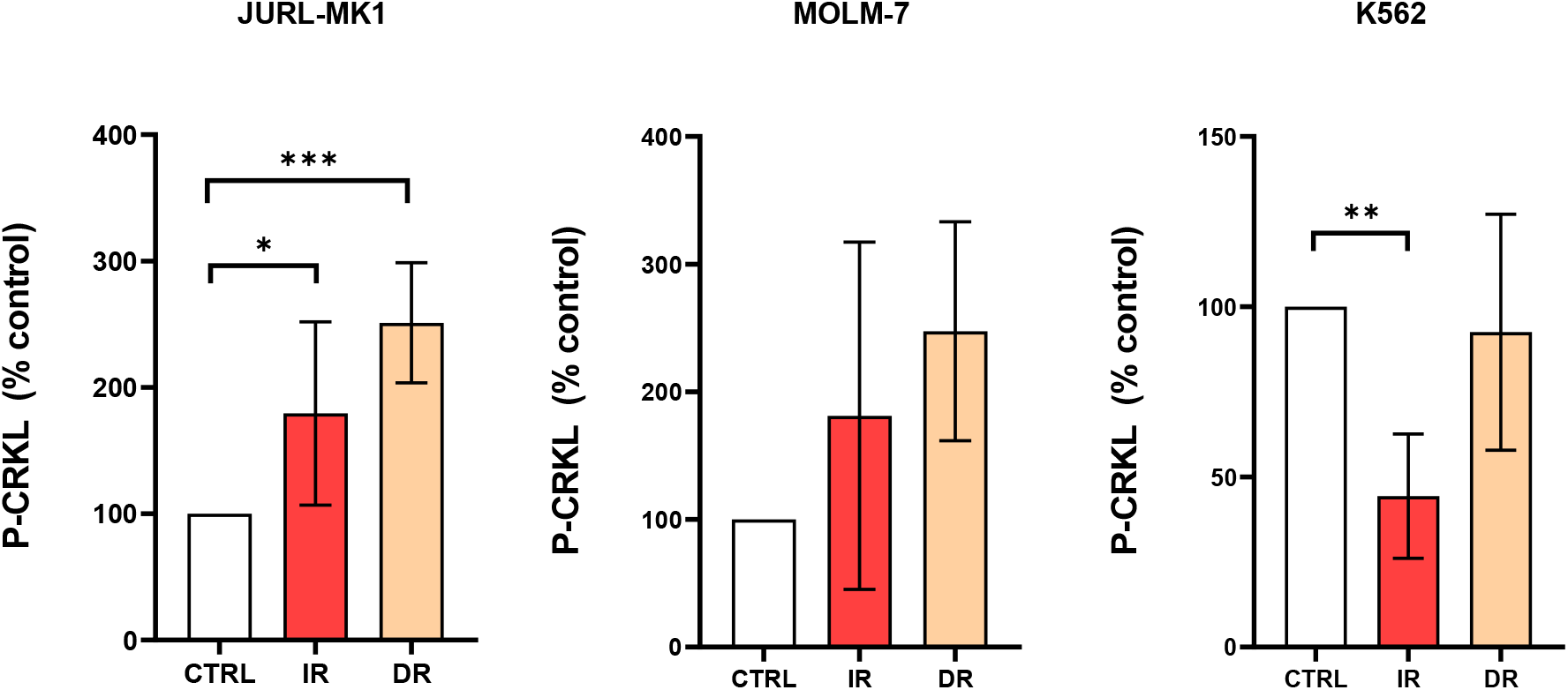
BCR-ABL activity of TKI-resistant cells. Relative densitometric graphs of phosphorylation of CRKL, surrogate marker of BCR-ABL activity. C—control, IR—imatinib resistant, DR—dasatinib resistant. Phosphorylation levels were normalised to β-actin and related to the corresponding controls. Means and standard deviation obtained from at least 3 biological replicates are shown. (***P< 0.001; **P<0.01; *P<0.05).

### Phosphorylation of CK2 substrates correlates with CK2 kinase protein levels in CML cell lines resistant to imatinib and/or dasatinib

To verify the results obtained from protein arrays (supplementary figure 1), we analyzed the level of HSP90β P-S226 and CK2 level in TKI-resistant cells (Figure 2, supplementary figure S4). The level of phosphorylated HSP90β-S226 was significantly increased in both JURL-MK1 derived, TKI-resistant sub-lines, compared to the sensitive counterparts. We therefore assessed the expression of CK2 subunits and evaluated CK2 activity using a specific antibody for the detection of CK2 phosphorylated substrates (Figure 2B).

**Figure 2.**
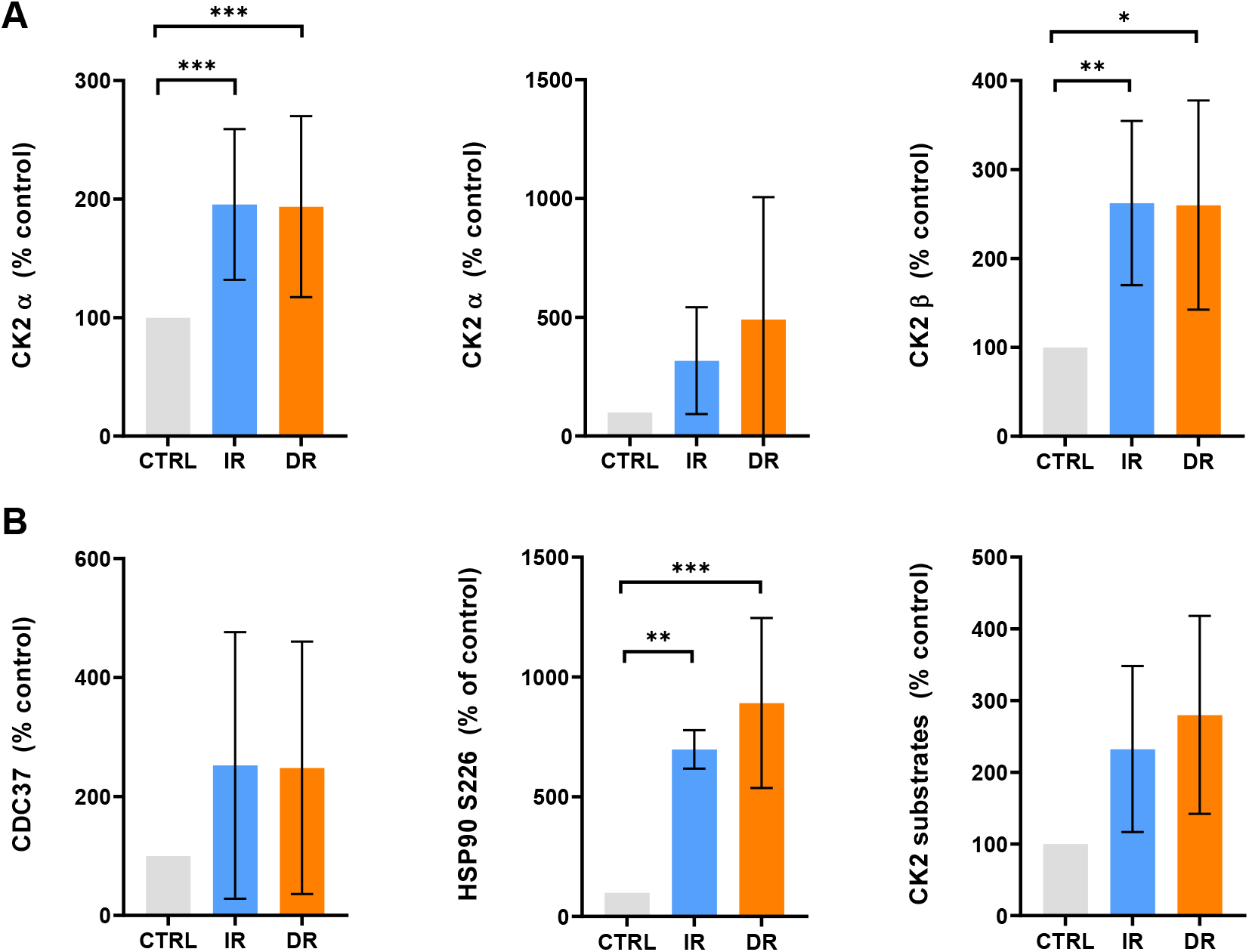
CK2 subunits and CK2 substrate phosphorylation in JURL-MK1 cells. Western blot analysis of JURL-MK1 cells and their resistant counterparts using antibodies against (A) CK2 subunits and (B) CK2 substrates CDC37, HSP90S226, and general CK2 consensus substrate sequence (pS/pT)DXE. Protein/phosphorylation levels were normalized to β-actin and related to the corresponding controls. Means and standard deviation obtained from at least 3 biological replicates are shown. (***P < 0.001; **P<0.01; *P<0.05).

While the CK2 α and β subunit levels were increased in both IR and DR JURL-MK1 cells (Figure 2A), the case could not be made for MOLM-7 resistants (Supplementary figure S4), and in K562 cells, the CK2 subunit expression was lower in resistant cells than in the parental cell line (Supplementary Figure S4B). In all cells, phosphorylation of CK2 substrates and CDC37 was correlated with the CK2 subunit levels (Figure 2B, Supplementary figure S4A, B).

Phosphorylation of the HSP90 co-chaperone CDC37 on S13 is known to be mediated by CK2 and can be used as a marker of CK2 activity (29). Given the fact that CDC37 protein was phosphorylated at S13 in all cell lines, we used it in subsequent experiments to assess CK2 inhibition.

### Effects of CK2 inhibition in TKI-resistant cells

To inhibit CK2, we used CX-4945 (Silmitasertib) as an inhibitor of choice. Based on the literature (30) and our own preliminary experience, 10 μM CX-4945 was used in all subsequent experiments.

### CK2 inhibition reduces cell proliferation and viability, and induces apoptosis in TKI-resistant cells

Proliferation ability of the cells was measured by Alamar Blue. Changes induced by TKIs and CX-4945 are shown in Figure 3A, B.

**Figure 3.**
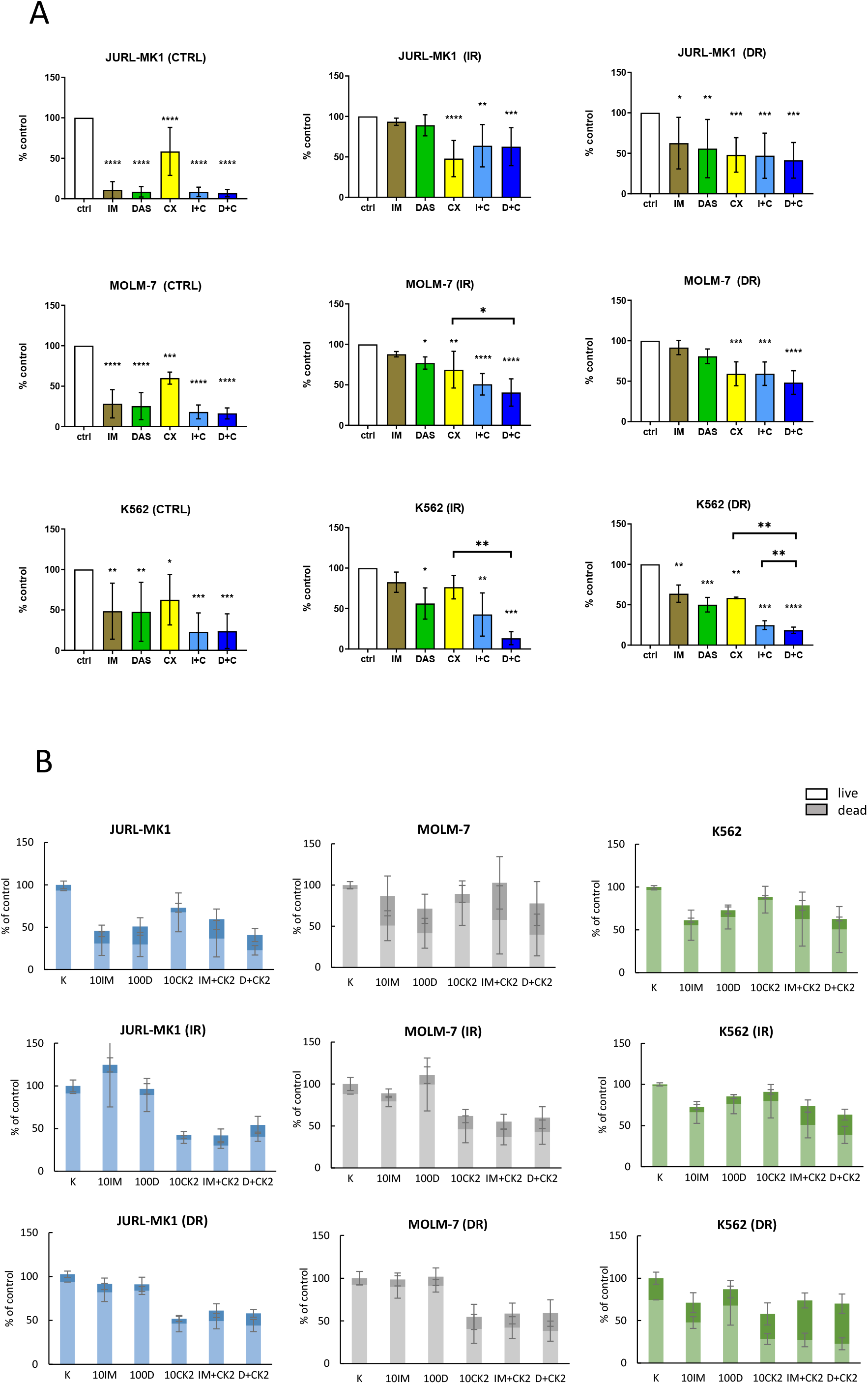
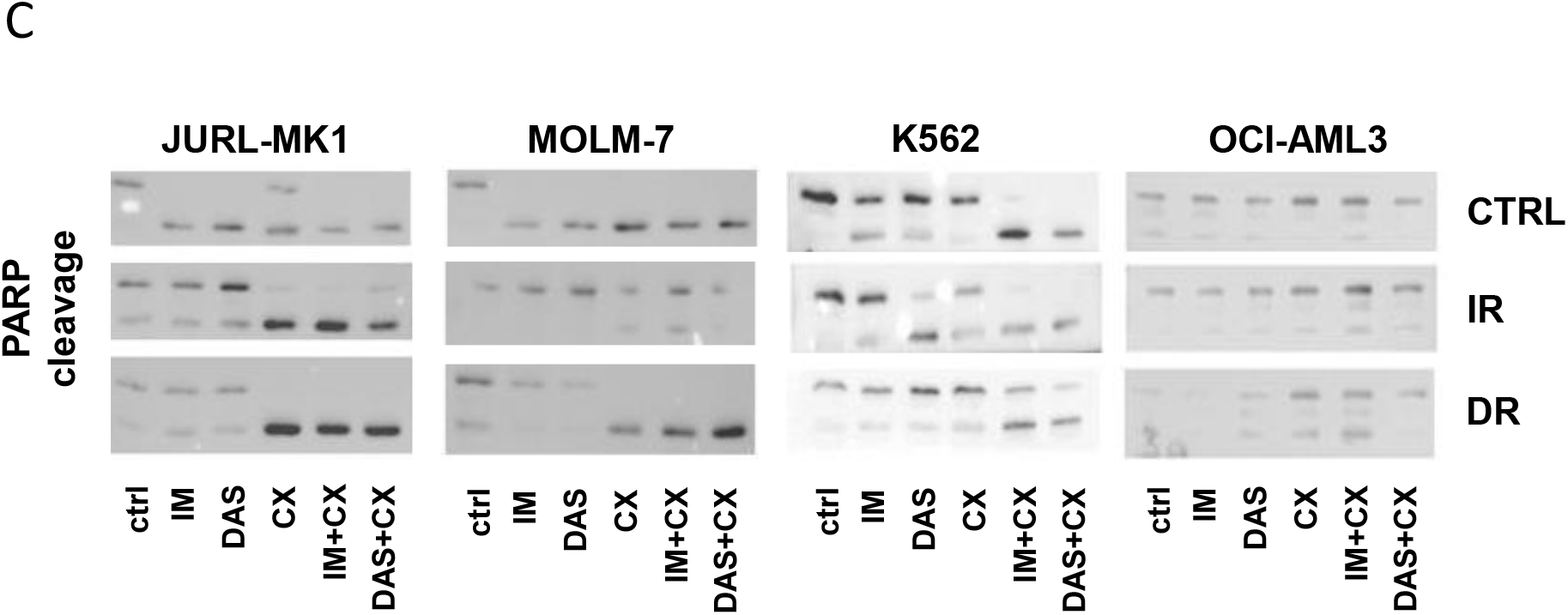
Effects of CX-4945 on cell proliferation, viability, and PARP cleavage. The cells were treated for 48 h with imatinib (IM; 10 μM), dasatinib (DAS; 100 nM), CX-4945 (CX; 10 μM), and their combinations. (A) Proliferation activity of the cells was assessed by the Alamar blue method. The data represents mean and SD of 3 – 6 separate experiments. Statistical significance was assessed by one-way ANOVA followed by a Dunnett’s multiple comparisons test. (B) Cell count and viability were evaluated by trypan blue staining. (C) Apoptosis was tested via cleaved PARP1 by western blots. Representative western blots are presented. Means and standard deviation obtained from at least 3 independent biological replicates are shown. ****P<0.0001, ***P < 0.001, **P<0.01, *P<0.05. C— control, IR—imatinib resistant, DR—dasatinib resistant.

While CX-4945 alone decreased cell proliferation in all parental CML cells (Figure 3A), the effect of TKIs alone was more pronounced. The effect of CX-4945 was not, however, affected by the newly acquired resistance, while the previously stronger effects of TKIs were largely diminished in the cells with acquired resistance. Importantly, with the exception of JURL-MK1 resistants, the combination of CX-4945 with TKIs had more profound effect on the resistant cells than either inhibitor alone, indicating potentiation. In BCR-ABL negative OCI-AML3, the effect of inhibitor combination was comparable to that of CX-4945 alone, and was not affected by acquired resistance of the OCI-AML3 cells (Supplementary figure 5A).

Similar results were observed for the cell count and dead cell fraction. While proven effective in JURL-MK1 and MOLM-7 sensitive cell lines, TKIs failed to exhibit an appreciable effect on the resistant cells (Figure 3B). In K562 cells, the effect of TKIs alone was similar for the sensitive and resistant cells. This correlated with the alamar blue proliferation assessment (Figure 3A). Once again, BCR-ABL negative OCI-AML3 cells were somewhat affected by CX-4945 treatment in terms of a slight growth slowing (Supplementary figure 5B), but CK2 inhibition did not induce an increase in the dead cell fraction in those cells.

In cases where TKIs or CX-4945 had taken effect on cell proliferation and viability, we were able to detect PARP cleavage, indicating apoptotic mechanism of the cell death (Figure 3C, Supplementary figure 5C).

### CK2 is regulated by BCR-ABL, and participates in the TKI resistance phenotype

There are opposing views on the issue of how CK2 and BCR-ABL regulate each other (31,32). We therefore aimed to evaluate the relationship of those kinases in our new cell models. To grasp this concept, we utilized the prominent substrates for CK2 and BCR-ABL, proteins CDC37 and CRKL, and assessed changes in their phosphorylation after treatment with inhibitors (Figure 4).

**Figure 4.**
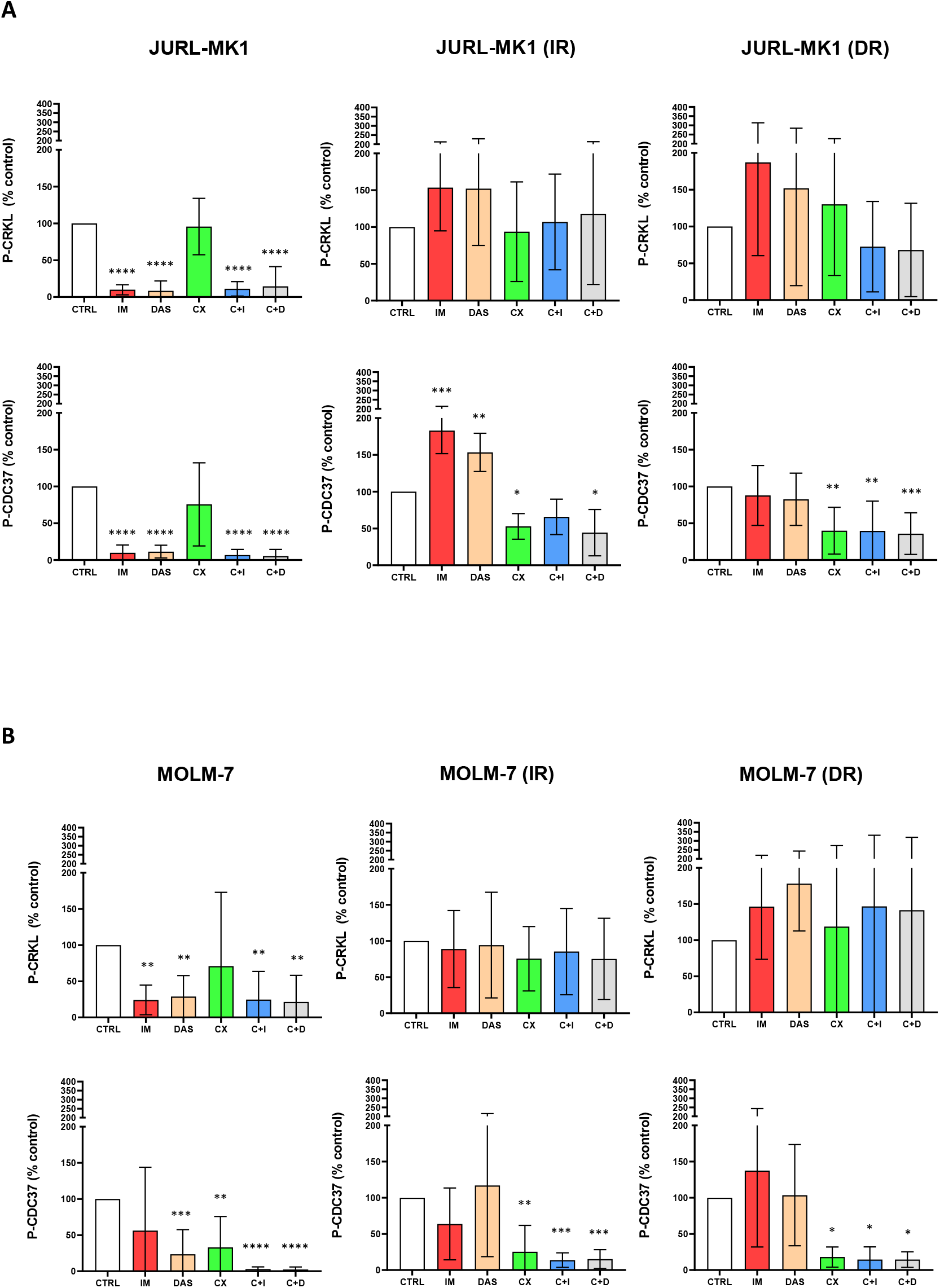
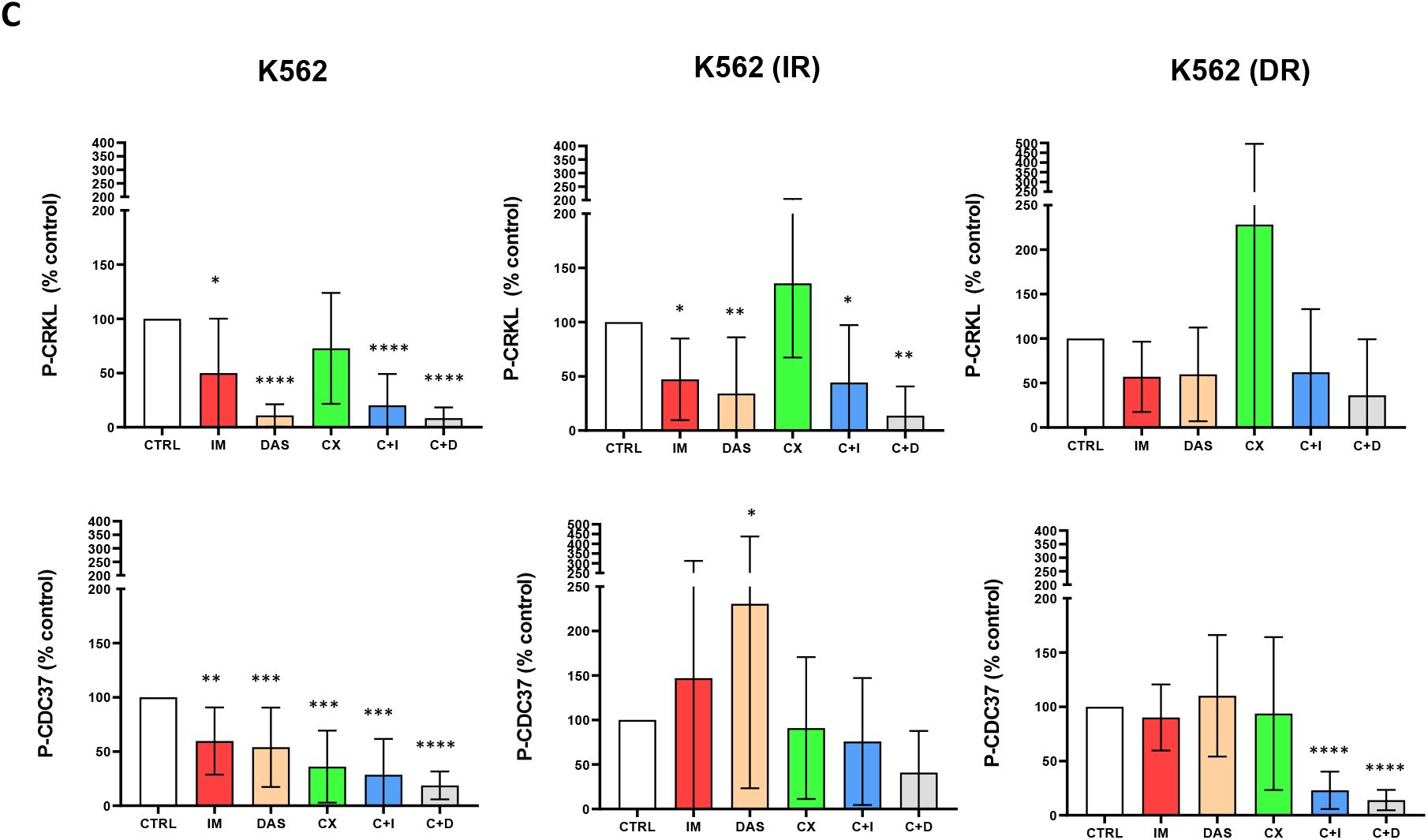
Effect of CX-4945 and TKIs on CK2 and BCR-ABL activity. The cells were treated for 48 h with imatinib (10 μM, IM), dasatinib (100 nM, DAS), CX-4945 (10 μM, CX), and their combinations. Quantification of western blot analyses of their effect to phosphorylation of CDC37 and CRKL are shown. Means and standard deviation obtained from at least 3 biological replicates. (A) JURL-MK1, (B) MOLM-7, (C) K562. ****P<0.0001, ***P < 0.001, **P<0.01, *P<0.05

According to the expectations, TKIs had significant effect on CRKL phosphorylation in sensitive cells, while CX-4945 did not significantly alter the P-CRKL levels, indicating that CK2 inhibition does not affect BCR-ABL activity. Phopshorylation of CDC37, on the other hand, was affected by the treatment with TKIs as well as with CX-4945, indicating that BCR-ABL activity is necessary for CK2 activation.

In resistant cells, CRKL phosphorylation in JURL-MK1 and MOLM-7 resistant cells remained largerly unchanged upon inhibitor treatment; in K562 cells, however, TKI treatment induced a slight reduction in CRKL levels. This may explain the slightly impaired reduction of proliferation and viability observed upon TKI treatment in the K562 resistants (see Figure 3A). Phosphorylation of CDC37 in the resistant cells was largely unaffected by TKIs, the exception being JURL-MK1_IR cells, where CDC37 phosphorylation was significantly higher in TKI-treated cells. The same trend was observed in K562_IR cells, but was not statistically significant. In JURL-MK1 and MOLM-7 resistants, CX-4945 reduced CDC37 phosphorylation alone and in combination with the TKIs; in K562 resistant cells, the effect of combined CK2 and BCR-ABL inhibition was more profound that CK2 inhibition alone. This was statistically significant for K562 DR cells.

### CK2 inhibition decreases viability and induces apoptosis of primary CML cells

To probe the potential of CK2 inhibition in primary CML cells, we collected samples from 2 CML patients without a satisfactory response to other TKIs (nilotinib and bosutinib), and without any mutation in the BCR-ABL kinase domain (for patient characteristics, see Table 3).

**Table 3.**
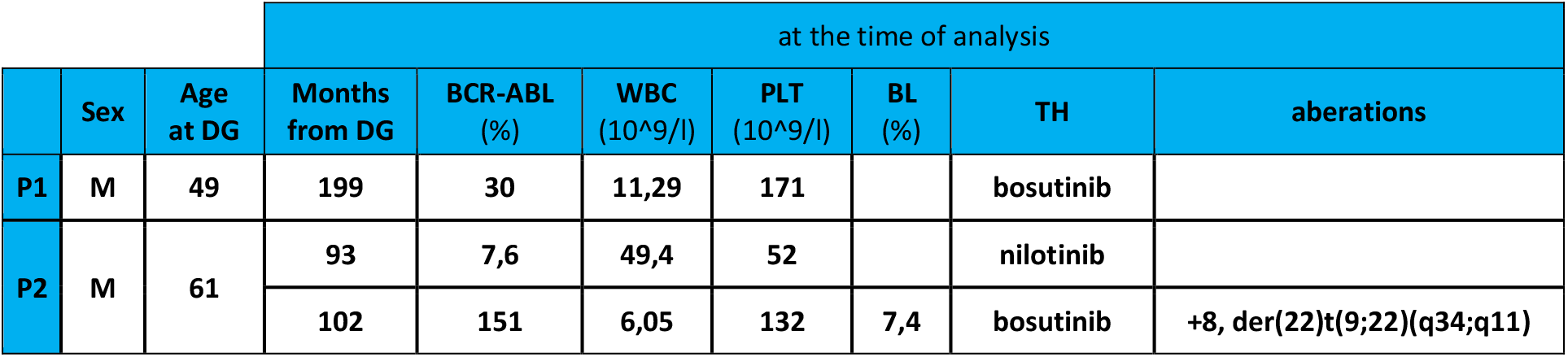
Patient characteristics. DG—diagnosis, WBC—total leukocyte count, PLT—platelet count, BL—blasts

We therefore incubated the cells with imatinib, dasatinib, and CX-4945 in the same conditions as we did previously with the cell lines. The patient differed in their (non)response to TKIs, patient 1 being more sensitive than patient 2 (Figure 5A). The sensitivity was somewhat unexpected due to the bosutinib clinical failure (refer to Table 3).

We further probed the potential effect of TKI / CX-4945 combination (Figure 5B). In patient no. 2 (insensitive to TKIs), compared with healthy donors, the combination of CX-4945 with TKIs highly decreased proliferation of the patient cells, while keeping only a mild effect on healthy donor samples.

**Figure 5.**
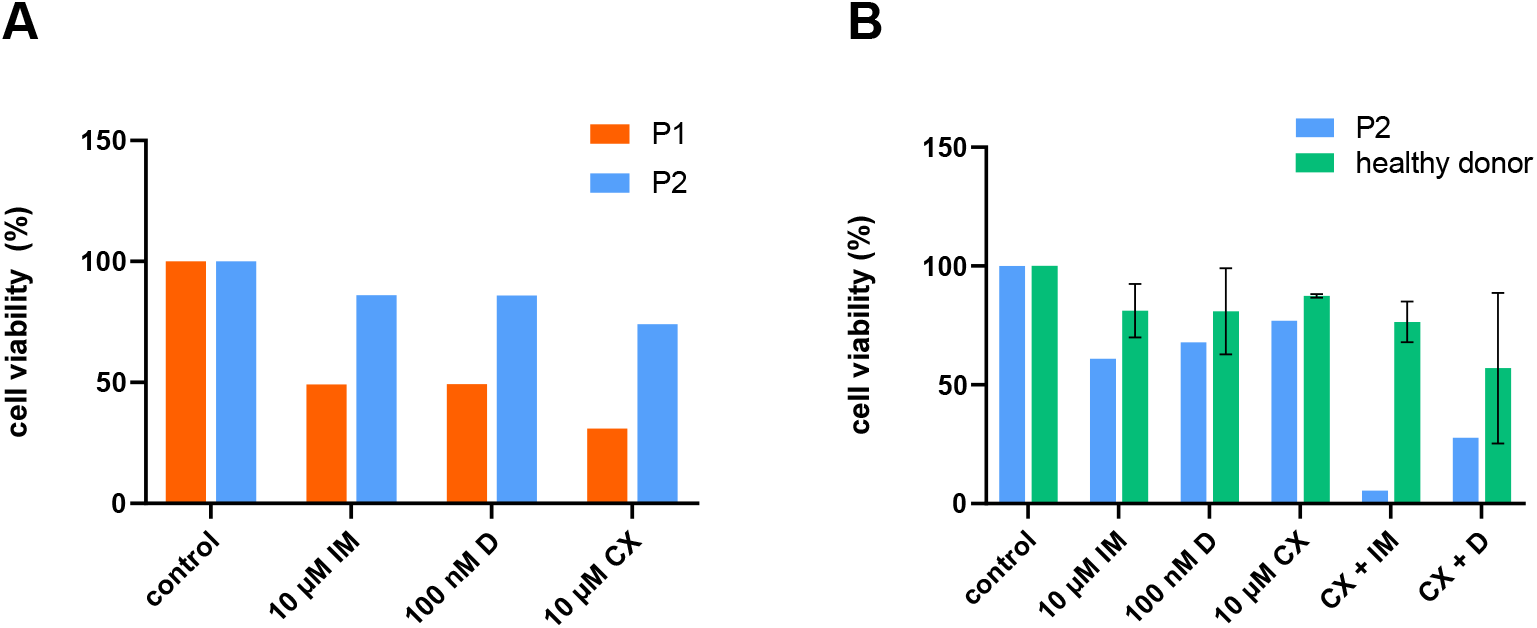
Effect of CK2 inhibition on primary cells from CML patients. (A) The cells collected from two CML patients non-responding to therapy were treated with imatinib (IM), dasatinib (D), or CX-4945 (CX) at indicated concentrations. Cell viability was assessed after 48 h incubation. (B) The cells from patient 2 (P2) and cells from healthy donors (HD; N=2) were incubated with the inhibitors as indicated. Viability after 48 h incubation was assessed.

## Discussion

Previously, we have shown that increased HSP90 levels correlate with negative outcomes in the CML course and response to therapy (33). Here, we found the HSP90β S226 phosphorylation increased in patients with poor response (either non-responding or relapsed) to TKI treatment (Supplementary figure 1A). Changes in HSP90β phosphorylation (both hyperphosphorylation and hypophosphorylation) are associated with TKI resistance in CML (34). In our case, the ratio of HSP90β P-S226 to total HSP90 was markedly increased even after one year of treatment in a patient not responding to therapy (Supplementary figure 1B).

HSP90 is a known substrate of CK2 (35–37). Given the known role of CK2 in several cancers, we hypothesized that CK2 plays a role in CML resistance to TKIs. Since the availability of patient samples is limited, we developed six new, CML-derived cell lines resistant to TKIs. All the cell lines designated as “resistant” were able to grow in media containing 2 μM imatinib or 2 nM dasatinib (Table 2, Supplementary Figure 2). The observed cross-resistance (insensitivity of the cells to another TKI than that used for resistance acquisition) confirms that the resistance mechanisms developed under the selective pressure of one inhibitor often provide resistance to other TKIs as well (38).

We characterized the cells as to their cytogenetic profile, BCR-ABL activity, and CK2-related protein content (Supplementary figures S3, S4, Figure 1, 2) to illustrate differences among these cell lines. Mutation of the BCR-ABL kinase domain is the main mechanism by which CML cells acquire resistance to TKIs, and in cases where BCR-ABL mutations do not develop, BCR-ABL gene amplification is usually present (39,40). Here, JURL-MK1 resistant cells developed the T315I mutation in the BCR-ABL kinase domain but showed no additional amplification of the BCR-ABL fusion gene, MOLM-7 and K562-derived resistants did not bear BCR-ABL mutation, but the BCR-ABL gene was amplified (Supplementary figure S3), in MOLM-7_R more than in K562_R cells. This leads us to the comparison of BCR-ABL acitivity in those cells (Figure 1), where in JURL-MK1 and MOLM-7 cells the phosporylation of a BCR-ABL activity surrogate marker, CRKL, was markedly increased. Interestingly, P-CRKL signal was significantly decreased in K562_IR cells compared to TKI-sensitive controls, despite the IR cells possessing additional (albeit non-amplified) BCR-ABL fusions in their genome (Supplementary figure S3). This is not unexpected, as others (41,42), also observed amplification of BCR-ABL gene without any KD mutation in imatinib-resistant cell lines.

The changes in expression of CK2 subunits in resistant cells compared to their sensitive counterparts was variable among cell lines (Figure 2, Supplementary figure S4) and in JURL-MK1 and K562 cells, the expression change in CK2 subunits correlated with the phosphorylation of CK2 substrates and CDC37, a surrogate marker for CK2 activity. In MOLM-7 cells, the expression of CK2 subunits did not change in the resistant cells, but we observed a trend of CK2 substrate phosphorylation increase. This may be explained, at least partially, by the interplay between CK2 and BCR-ABL (Figure 4).

In CML, CK2 interacts with BCR-ABL (35,36), and imatinib-resistant cells seem to be sensitive to CK2 inhibitor CX-4945 (21,25). This is also reflected in our cell models (Figure 3A,B). Additionally to that, we show that the same effect can be observed in dasatinib-resistant cells. Targeting CK2 with CX-4945 resulted in a decrease in cell viability and subsequent induction of cell death in most of the CML cells used in our study (Figure 3), even though BCR-ABL signaling remained active in all cells after CK2 inhibition (Figure 4). Additionally, targeting BCR-ABL with imatinib or dasatinib affected CK2 activity in sensitive cells, but not in resistant cells (Figure 4). While in cells with high BCR-ABL activity (i.e. JURL-MK1 and MOLM-7 resistants, see Figure 1) the activity of CK2 correlated with that of BCR-ABL, in K562 resistants the activity of CK2 remained the same (DR) or even increased (IR) despite BCR-ABL being suppressed. This suggests that in resistant cells, CK2 activity is dependent on that of BCR-ABL; in resistant cells, these mechanisms may change, and uncoupling of BCR-ABL and CK2 mutual regulation may be a part of the resistant phenotype. This is supported by the K562 resistant cells, in which the “uncoupling” is fairly obviously lost, and simultaneously, these cell’s resistance phenotype appears to be the least dependent on BCR-ABL compared to JURL-MK1 and MOLM-7 resistants. Other candidate molecules to mediate K562 resistance are Src family kinases (SFK), which are recognized as important players in TKI resistance (43,44), and are know to regulate and be regulated by CK2 (45,46).

Importantly, even in resistant cells, the combination of TKI and CX-4945 was in all cases the most effective on viability and proliferation parameters (Figure 3), irrespectively of the BCR-ABL activity (Figure 4). Together, this indicates that while active BCR-ABL kinase is not affected by the CK2 activity decrease, the effects on cell proliferation and the increase of apoptotic parameters (Figure 3C) support the CK2 “addiction” hypothesis (17) in CML cells as well. CK2 inhibition therefore may help to overcome CML resistance to TKIs.

As we and others have previously shown (47–49), results of *in vitro/ex vivo* cultivation of isolated patient leukocytes with TKIs correlate well with therapy results and are of a great prognostic value. Similarly, in vitro experiments with CK2 inhibitors could predict the role of CK2 in resistance to TKI and thus a possibility to target CK2 kinase. CK2 inhibition did not affect leukocytes from healthy donors (Figure 7), which corresponds to previous reports (29). On the other hand, CK2 inhibition alone or in combination with TKI decreased the metabolic activity of primary cells of CML patients with resistance to various TKI (Figure 7C).

In conclusion, we present a study of the effects of CK2 inhibition on TKI-resistant CML cells. Using six originally developed cell lines resistant to tyrosine kinase inhibitors as well as primary patients’ samples, we showed that CK2 inhibition and/or its combination with TKIs is capable of inducing cell death in TKI-resistant cells. This is true even for cells bearing the gatekeeper T315I mutation. Importantly, BCR-ABL negative cell line HL60 and primary cells from healthy donors remain unaffected by such treatment.

## Material and Methods

### Patient samples and cell cultures

Leukocytes from CML patients and healthy donors were isolated as described previously (33,49). Responses to therapy were defined according to European Leukemia Net (ELN) classification (22). Samples were obtained with the agreement of the Ethics Committee of the Institute of Hematology and Blood Transfusion (Prague, Czech Republic) according to the Declaration of Helsinki. Written informed consent was obtained from all patients.

Cell lines were purchased from DSMZ (Braunschweig, Germany – OCI-AML3, JURL-MK1) or ECACC (Salisbury, UK – HL-60, K562). The cell line MOLM-7 (not commercially available) was obtained from Dr J. Minowada. From all used cell lines, sub-lines resistant either to 2 μM imatinib or 2 nM dasatinib were raised according to the protocol by Mahon et al. (50). Briefly, resistant cells were developed by prolonged exposures to gradually increasing concentrations of imatinib and dasatinib, starting from 1 nM to 2 μM (imatinib) and 1 pM to 2 nM (dasatinib). Parental cell lines were cultivated in parallel without inhibitors. All cells were incubated at 37 °C in RPMI medium with 10% fetal bovine serum and antibiotics.

### Chemicals and antibodies

Imatinib mesylate, nilotinib, dasatinib, and CX-4945 were purchased from Santa Cruz Biotechnology, Inc. All inhibitors were dissolved as 10 mM stock solution in DMSO (Sigma-Aldrich Inc., Missouri, USA) and stored at −20 °C. Protease inhibitor cocktails and Phosphatase inhibitor cocktails were purchased from Calbiochem and SIGMA-ALDRICH, respectively. Non-conjugated primary antibodies casein kinase IIα (D-10 and E-7), casein kinase IIα′ Antibody (E-7), casein kinase IIβ Antibody (6D5), and Actin (anti-actin antibody clone AC-15, SIGMA-ALDRICH) or beta-tubulin (β-Tubulin (9F3) were purchased from SantaCruz Biotechnology (CA, USA). Anti-HSP90 beta (phospho S226) antibody was purchased from Abcam (UK). Phospho-CK2 Substrate [(pS/pT)DXE] MultiMab™ Rabbit mAb, Phospho-CDC37 (Ser13) (D8P8F) Rabbit mAb, Phospho-CrkL (Tyr207) Rabbit mAb, Cleaved PARP (Asp214) (D64E10) XP, and HRP conjugated secondary antibodies were purchased from Cell Signaling Technology (Danvers, MA, USA).

### Cell viability and proliferation

Cell viability was assessed by Trypan blue staining (according to the manufacturer’s protocol); cell metabolic activity was evaluated using the AlamarBlue® assay (Invitrogen), following the manufacturer′s instructions. Briefly, 100 μL of cell suspension was transferred to a 96-well plate and 10 μL of AlamarBlue reagent was added to each well. After incubation at 37 °C for 1 hour, fluorescence intensity was measured on microplate reader BMG FLUOstar Galaxy (MTX Lab Systems, Inc., VA, USA). Statistical analysis was performed using GraphPad Prism 8.2.

### Immunoblotting

Western blot analyses were performed as described previously 68. Briefly, cells were harvested, lysed in Laemmli buffer, and boiled for 10 minutes. The samples were resolved on SDS polyacrylamide gel (10 %), transferred to PVDF membrane, and incubated with the appropriate antibodies. Protein bands were detected by chemiluminescence (SuperSignal West Dura Extended Duration Substrate -Thermo Fisher Scientific, USA) and scanned using a G:BOX imager (Syngene Europe). Densitometric quantification was performed using Gene Tools product version: 4.3.8.0 (Syngene Europe). All analyses were repeated 2-5 times.

### BCR-ABL transcript variants analysis

#### RT-PCR

Total RNA was extracted by the NucleoSpin® RNA Plus Kit (Macherey-Nagel, Germany) according to the appropriate protocol. The cDNA synthesis was prepared using oligoT primers and SuperScript II transcriptase (Invitrogen/Life Technologies, USA). The cDNA of CML leukemic cell lines and control cell lines was used during the development. PCR was performed using a CFX96 Touch Real-Time PCR Detection Systems (Bio-Rad). Primer sequence for a2b2 (e13b2) and a2b3 (e14b2) transcript variants:

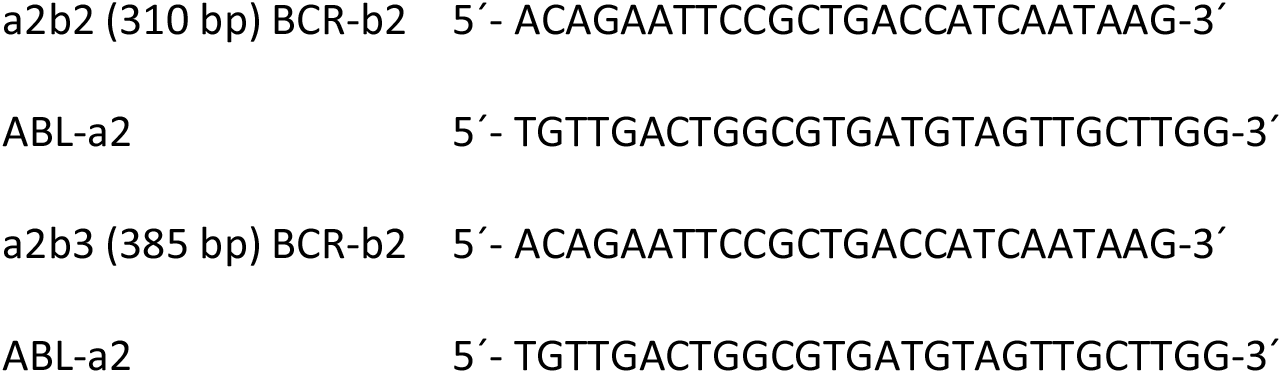

#### Nested PCR

Long-range nested RT-PCR analysis was performed to amplify BCR-ABL cDNA. For this purpose the sequences of primers used for RT PCR were as follows:

Step 1:

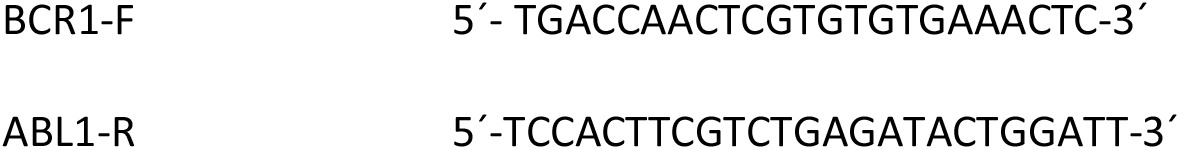

Step 2:

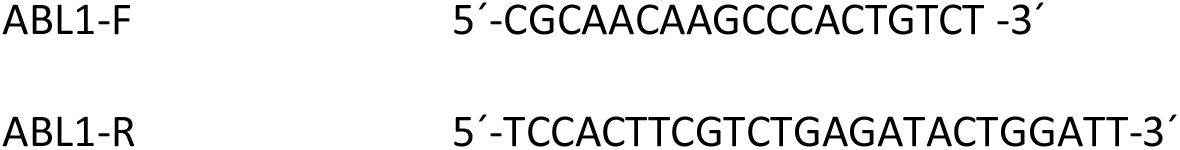

Step 1: initial denaturation at 95 °C / 2 min, 40 cycles at 95 °C / 1 min, 60 °C / 1 min, and 72 °C / 3 min. Final 10-min extension step at 72 °C. Step 2: initial denaturation at 95 °C / 2 min, 50 cycles at 95 °C / 1 min, 60 °C / 1 min, and 72 °C / 3 min. Final 10-min extension step at 72 °C.

### Mutational analysis

PCR products were separated on 2% agarose gel containing MIDORIGreen Advance. Appropriate bands were cut out and purified with the QIAquick Gel Extraction kit (Qiagen). Sanger sequencing was performed on ABI PRISM 3500 Genetic Analyzer using Big Dye Terminator 3.1 kit (Applied Biosystems). The resulting sequences were analyzed in Chromas 2.31 program and tools/program of Blast was used for alignment of sequences.

### Cytogenetics

Cells were treated with demecolcin for 1,5 hours. Harvesting and preparation of slides were performed according to standard cytogenetic procedures. Cells were stored at −20 °C in methanol-glacial acetic acid (3:1). For cytogenetic analyses, cell suspensions were dropped on microscopic slides and air-dried. Fluorescence in situ hybridization (FISH) in combination with multicolor fluorescence in situ hybridization (mFISH) were used to characterize the chromosomes. Analyses were performed using commercially available probes BCR/ABL Vysis LSI BCR/ABL Dual Color, Dual Fusion Translocation Probe (Abbott Vysis, USA), and 24 XCyte mFISH Kit (MetaSystems, Germany). All available mitoses for both probes and two hundred nuclei for the LSI BCR/ABL probe were analyzed using an AxioImager Z1 fluorescence microscope (Carl Zeiss, Germany) and the Isis computer analysis system (MetaSystems, Germany). Findings were described according to ISCN 2016 (51).

### Phosphorylation antibody microarray

The Phospho Explorer Antibody Array PEX100 containing 1,318 well-characterized site-specific antibodies was purchased from Full Moon Biosystems (Sunnyvale, CA, USA). The isolated whole fraction of leukocytes was processed according to the manufacturer’s instructions. Briefly: cells were lysed, labeled by Biotin, placed to glass array, and incubated overnight at 4 °C. Cy3-Streptavidin was used for protein visualization, signal to noise ratio was measured on GenePix® Microarray Scanner (4000A; Molecular Devices, USA).

### Statistical analysis

All statistical analysis was performed using GraphPad Prism 8.2 software (GraphPad Software, Inc.). Analyses were assessed by repeated-measures ANOVA, Dunnett’s multiple comparisons test was used. Statistically significant results were obtained in independent biological replicates. P < 0.05 was considered statistically significant. Experiments were repeated at least three times. All data are presented as the mean ± standard deviation.

## Authors contribution

OM: experiments, data analyses and manuscript drafting

DM: experiments, data analyses

TML: conceptualization and design, performance of experiments

AO: performance of experiments, manuscript drafting

KC: performance of mutation analyses

SR: performance of cytogenetic experiments, data analyses and manuscript drafting

JB: performance of cytogenetic experiments, data analyses and manuscript drafting

HK: provision of patient samples, clinical data evaluation

MZ: conceptualization & design, performance of experiments, data analyses and manuscript drafting. All authors criticaly revised the manuscript. The authors declare no relevant conflict of interest.

