## Supplementary data for "Inhibition of Casein Kinase 2 induces cell death in chronic myelogenous leukemia cells resistant to tyrosine kinase inhibitors"

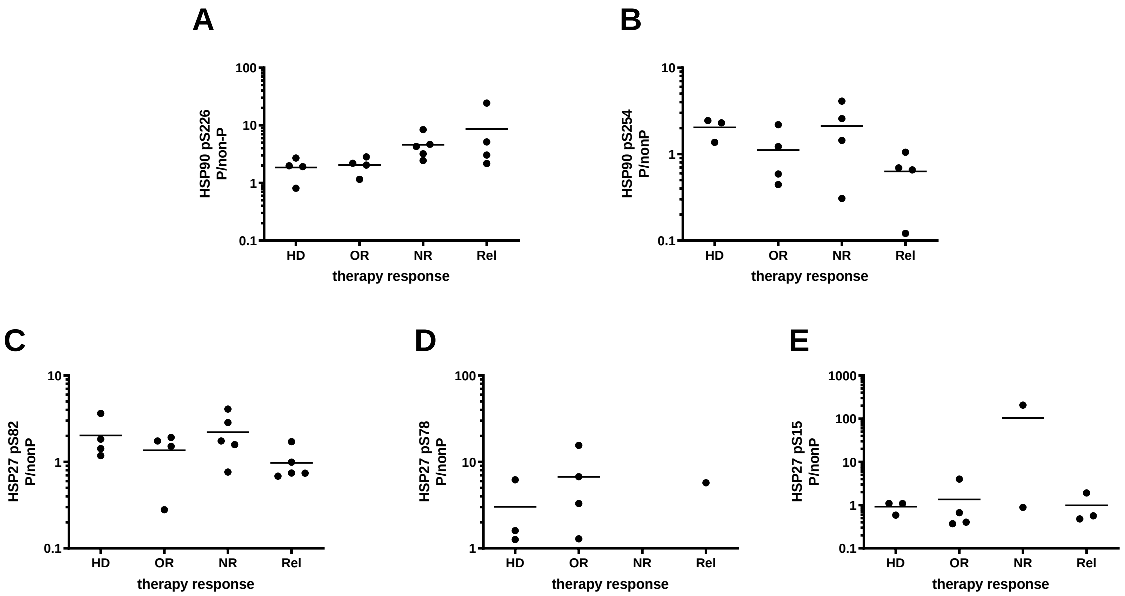


**Supplementary figure S1. HSP90 serine 226 phosphorylation levels in CML patients with differing responses to therapy.** Protein-antibody array analysis of 14 samples from patients with a different response to therapy. The ratios of phosphorylated and non-phosphorylated indicated serine residues are given. Therapy response: HD — healthy donor, OR — optimal response, NR — nonresponding, Rel — relapsed.

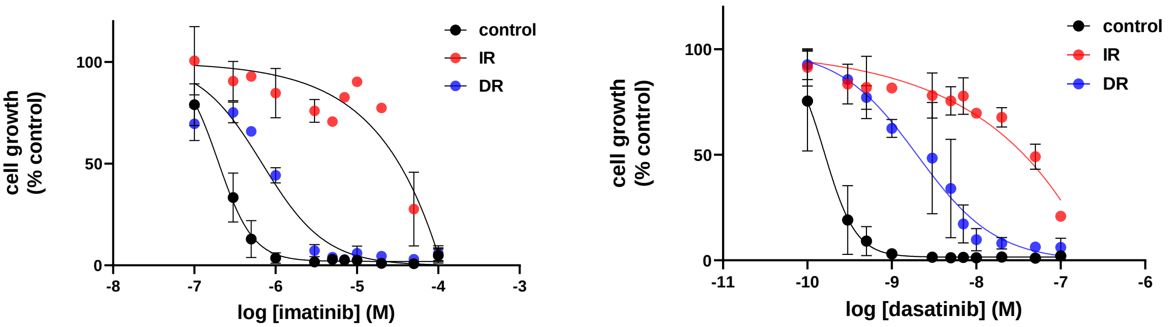


**Supplementary figure S2. Representative dose-response curves of JURL-MK1 cells.** JURL-MK1 cells and their resistant sub-lines were incubated with 0-100 µM imatinib or 0-100 nM dasatinib, as indicated. Graphs are means from 2-3 independent experiments, bars = s.d. IR — imatinib-resistant cells, DR — dasatinib-resistant cells.

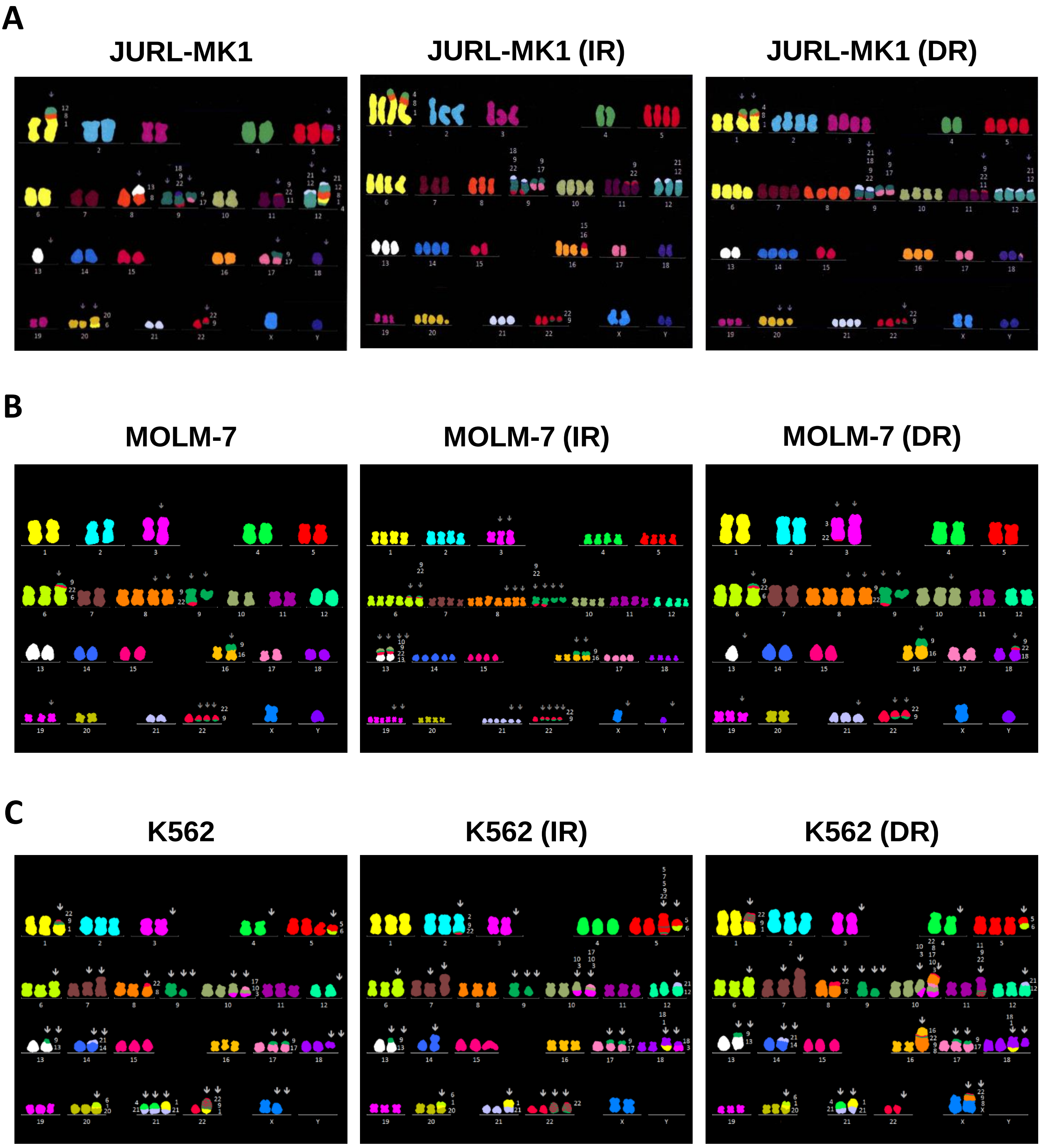


**Supplementary figure S3. Cytogenetic analysis of TKI-resistant cells.** Multicolor fluorescence in situ hybridization (mFISH) were used to characterize the karyotype of all CML-derived cells and their IR and DR sublines. (A) JURL-MK1, (B) MOLM-7, (C) K562. IR—imatinib resistant, DR—dasatinib resistant.

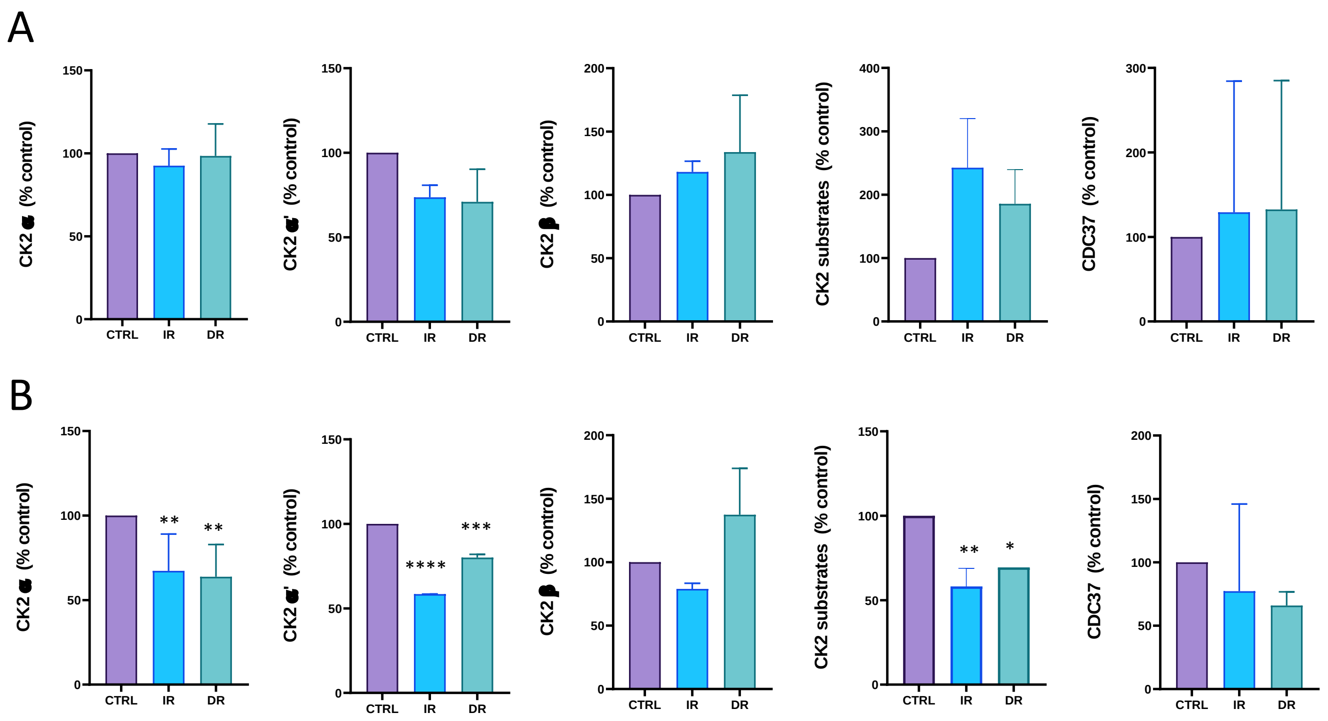


**Supplementary figure S4. Protein levels of CK2 subunits and its substrates phosphorylation in cell lines MOLM-7 (A) and K562 (B) and resistant their sub-lines.**
Densitometric graphs were calculated from western blots with appropriate antibodies. Phosphorylation and protein levels were normalised to β-actin and related to control. Means and standard deviation obtained from at least 3 biological replicates are shown. (***P < 0.001, **P<0.01, *P<0.1).

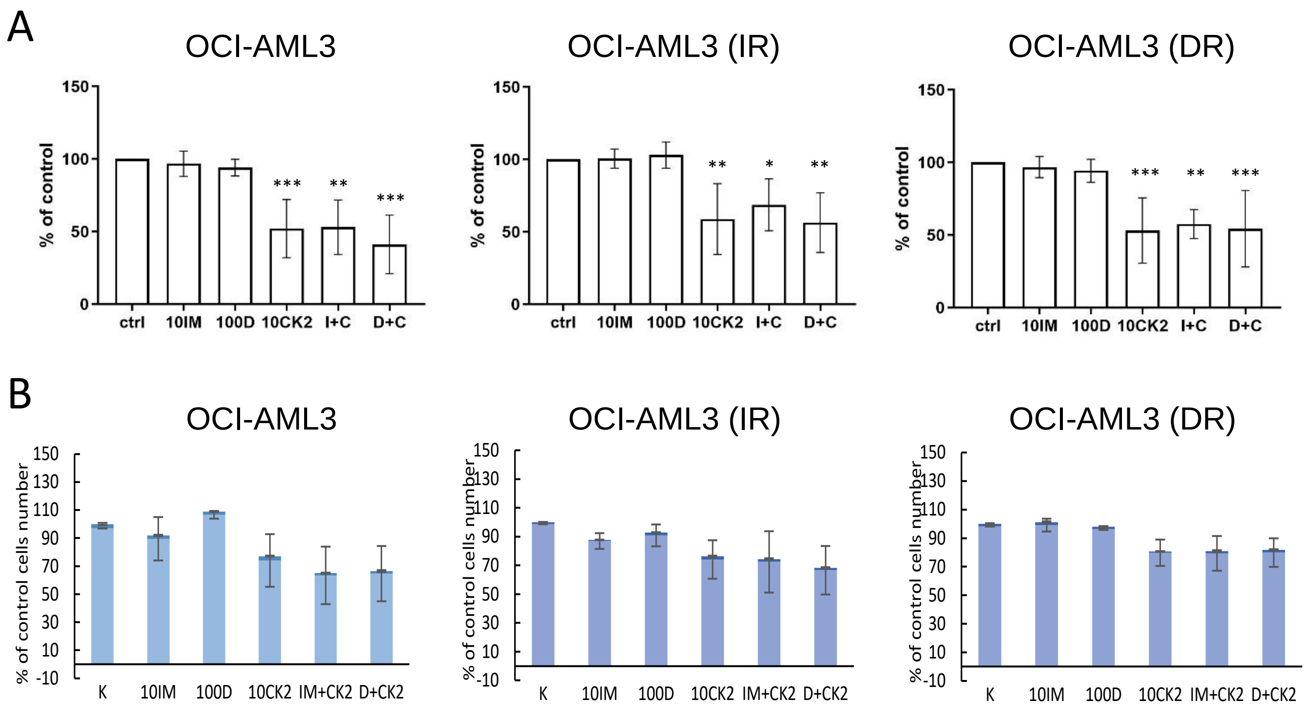


**Supplementary Figure 5. Effects of CX-4945 on cell proliferation and viability BCR-ABL negative OCI-AML3 cells.** The cells were treated for 48 h with imatinib (IM, 10µM), dasatinib (D, 100nM), CX-4945 (CK, 10µM), and their combinations. (A) Proliferation/cell activity of OCI-AML3 cells was assessed by the Alamar blue method and related to the control. The data represents mean with SD of 3 – 6 separate experiments, statistical significance was assessed by one-way ANOVA followed by a Dunnett's multiple comparisons test (B) Cell numbers and viability were evaluated by trypan blue staining. Light bars—live cells, dark bars—dead cell fraction.

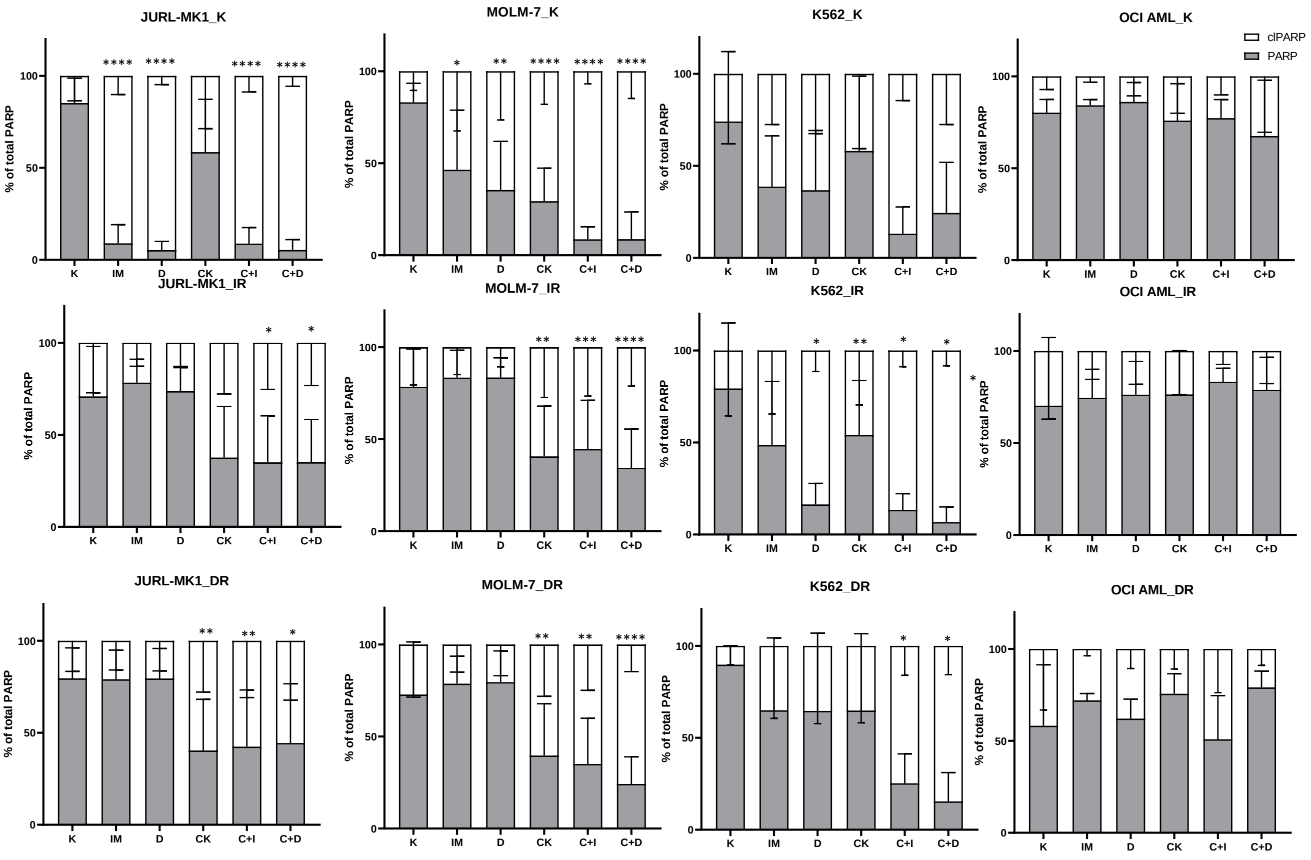


**Supplementary Figure 6. Effects of CX-4945 on the PARP cleavage.**

Densitometric evaluation of PARP/clPARP. Means and standard deviation obtained from at least 5 experiments (3 biological replicates, each at least 2 times analysed in western blot) are shown.

****P<0.0001, ***P < 0.001, **P<0.01, *P<0.05. C – control, IR – imatinib-resistant, DR – dasatinib-resistant.
